# Human bone tissue-derived ECM hydrogels: Controlling physicochemical, biochemical, and biological properties through processing parameters

**DOI:** 10.1101/2023.08.03.551765

**Authors:** Yang-Hee Kim, Gianluca Cidonio, Janos M. Kanczler, Richard OC Oreffo, Jonathan I Dawson

## Abstract

Decellularized tissues offer significant potential as biological materials for tissue regeneration due to their ability to preserve the complex compositions and architecture of the native extracellular matrix (ECM). While the use of decellularized ECM hydrogels from bovine and porcine bone tissues has been extensively studied, the evaluation and derivation of decellularized matrices from human bone tissue remain largely unexplored.

The objective of this study was to investigate how the physiochemical and biological properties of ECM hydrogels derived from human bone ECM could be controlled by manipulating bone powder size (45-250 μm, 250-1000 μm, and 1000-2000 μm) and ECM composition through modulation of digestion time (3, 5, and 7 days).

The current studies demonstrate that a reduction in material bone powder size and an increase in ECM digestion time resulted in enhanced protein concentrations in the ECM hydrogels, accompanied by the presence of a diverse array of proteins. Furthermore, these adjustments in the physicochemical properties generated improved gelation strength of the hydrogels. The evaluation of human bone marrow-derived stromal cells (HBMSCs) cultured on ECM hydrogels derived from 45-250 μm bone powder, treated for 7 days, demonstrated enhanced osteogenic differentiation compared to hydrogels derived from both larger bone powders and collagen gels.

In conclusion, this study highlights the significant promise of human bone ECM hydrogels as biologically active materials for bone regeneration. The ability to manipulate digestion time and bone powder size enables the generation of hydrogels with enhanced release of ECM proteins and appropriate gelation and rheological properties, offering new opportunities for application in bone tissue engineering.

## 1. Introduction

Tissue engineering offers significant promise for the repair of damaged tissues using cells, bioactive molecules, and biomaterials. Biomaterials play a crucial role providing a microenvironment and, typically, delivery of molecules that can promote the functions of the requisite cells within the regenerating tissues. To achieve this, numerous strategies have focused on developing and designing biomaterials that can mimic the structure and composition of the native extracellular matrix (ECM) ^[1, 2]^. Natural polymer-based hydrogels, such as collagen, alginate, chitosan, hyaluronic acid, and gelatin, display similar components as ECM and can create fibrous and porous microstructures, regulating cell behavior including attachment, proliferation, and differentiation ^[3, 4]^. However, reproducing the complex mixture of structural and functional proteins in ECM, including glycosaminoglycans (GAGs), collagen, proteoglycans, and growth factors, remains, to date, challenging.

Decellularization of tissues has emerged as a promising approach for the generation and subsequent application of biological materials. Following the removal of the cellular components from native tissues, the complex compositions and architecture of the ECM can be retained, leaving a decellularized ECM ^[5, 6]^. Furthermore, decellularization can reduce immunogenicity reducing associated issues around inflammatory responses and disease transmission. Indeed, decellularized ECM has been shown to advance cell behavior and constructive tissue remodeling responses ^[7-9]^. Decellularised matrices have found application in various clinical products, including three-dimension implantable scaffolds, 2D sheets, and powders ^[6, 10]^. In recent decades, decellularised ECMs have been formulated as hydrogels ^[11]^. ECM hydrogels are primarily composed of proteins solubilized or digested from decellularised ECM, and these proteins with the capacity to form a gel at physiologic temperature and pH. As a consequence, such hydrogels can be applied at a defect site through injection providing a more homogenous concentration of material in contrast to the use of ECM powders ^[12]^. The most commonly used method for generating an ECM hydrogel is through pepsin-mediated solubilization of decellularised ECM powders. Pepsin, an enzyme, can solubilize a range of proteins from decellularised ECM powders. A number of studies have reported that ECM protein-based hydrogels can retain the inherent bioactivity of native ECM, leading to improvements in cell proliferation, differentiation, and tissue repair ^[13, 14]^ and, naturally, implementation in within tissue engineering and regenerative medicine ^[5, 11]^. To date, there have been a paucity of publications harnessing decellularized ECM hydrogels derived from bovine and porcine bone tissues for bone tissue regeneration. These hydrogels have demonstrated the ability to enhance cell function and promote bone regeneration ^[15, 16]^. In the preparation of porcine bone-derived ECM hydrogels, demineralized and decellularized bone powders (<40 μm) were treated in pepsin solution for 1 hour ^[17]^. Similar, ECM hydrogels derived from bovine bone powders were fabricated after 96 hours of digestion in a pepsin solution ^[15]^. However, to date, the effect of tissue size and digestion time on protein concentration, gelation, rheological and biological properties of ECM hydrogels remains unknown. Critically, there is no study on the fabrication of hydrogels derived from human bone tissue ECM. The current studies examined the influence of demineralized and decellularised human bone powder size and digestion time on the physicochemical and biological properties of human bone ECM hydrogels for potential application in bone tissue repair. Human trabecular bone tissues were obtained from human femoral heads following orthopedic surgery, and demineralized human bone powders of varying sizes (45 – 250 μm, 250 – 1000 μm, and 1000 – 2000 μm) were prepared. The powders were subjected to pepsin treatment for 3, 5, or 7 days. We evaluated the gelation and rheological properties of the resulting ECM hydrogels based on the powder size and digestion time. Hydrogel microstructure was examined using scanning electron microscopy and micro-computed tomography. In addition, the efficiency of protein digestion was assessed by measuring cellular and protein concentrations. The proteomics of the derived hydrogels were evaluated using mass spectrometry and human bone marrow-derived stromal stem cell attachment, spreading, migration, proliferation, and differentiation examined on the ECM hydrogels. This study set out to examine the impact of powder size and digestion time on the properties of ECM hydrogels derived from human bone tissue, to offer insights on the ECM physicochemical properties for the development of effective biomaterials for bone repair applications.

## 2. Results and Discussion

In light of the growing interest in replicating the properties of the native extracellular matrix (ECM) in the fields of biomaterials and tissue regenerative medicine, our objective was to develop hydrogels derived from human bone ECM for orthopaedic reparative applications. In this study, we proposed the hypothesis that the size of ECM bone powder particles and the duration of ECM digestion are crucial in determining the quality and quantity of proteins derived from the ECM, consequently influencing the rheological and physicochemical properties of the resulting hydrogel. This study investigated the potential of human bone ECM hydrogels to promote bone tissue repair through examination of human skeletal cell population function.

### 2.1. Generation of porous and fibrous hydrogel microstructures

ECM hydrogels derived from demineralised and decellularised human bone were successfully fabricated (Figure 1). To investigate the effect of bone powder size and digestion time, we prepared various types of human bone ECM hydrogels (Table 1) and examined the cross-section and 3- dimensional structures of the hydrogels. All the hydrogels examined exhibited a porous and fibrous structure (Figure 2a and Supplementary Figure 1). The fibers were widely distributed throughout the pores, which were interconnected (Figure 2a and b). The pore area percentage, calculated based on SEM images using ImageJ, did not display any significant differences between the hydrogels (51.8 ± 4.3 %) (Figure 2c).

**Figure 1.**
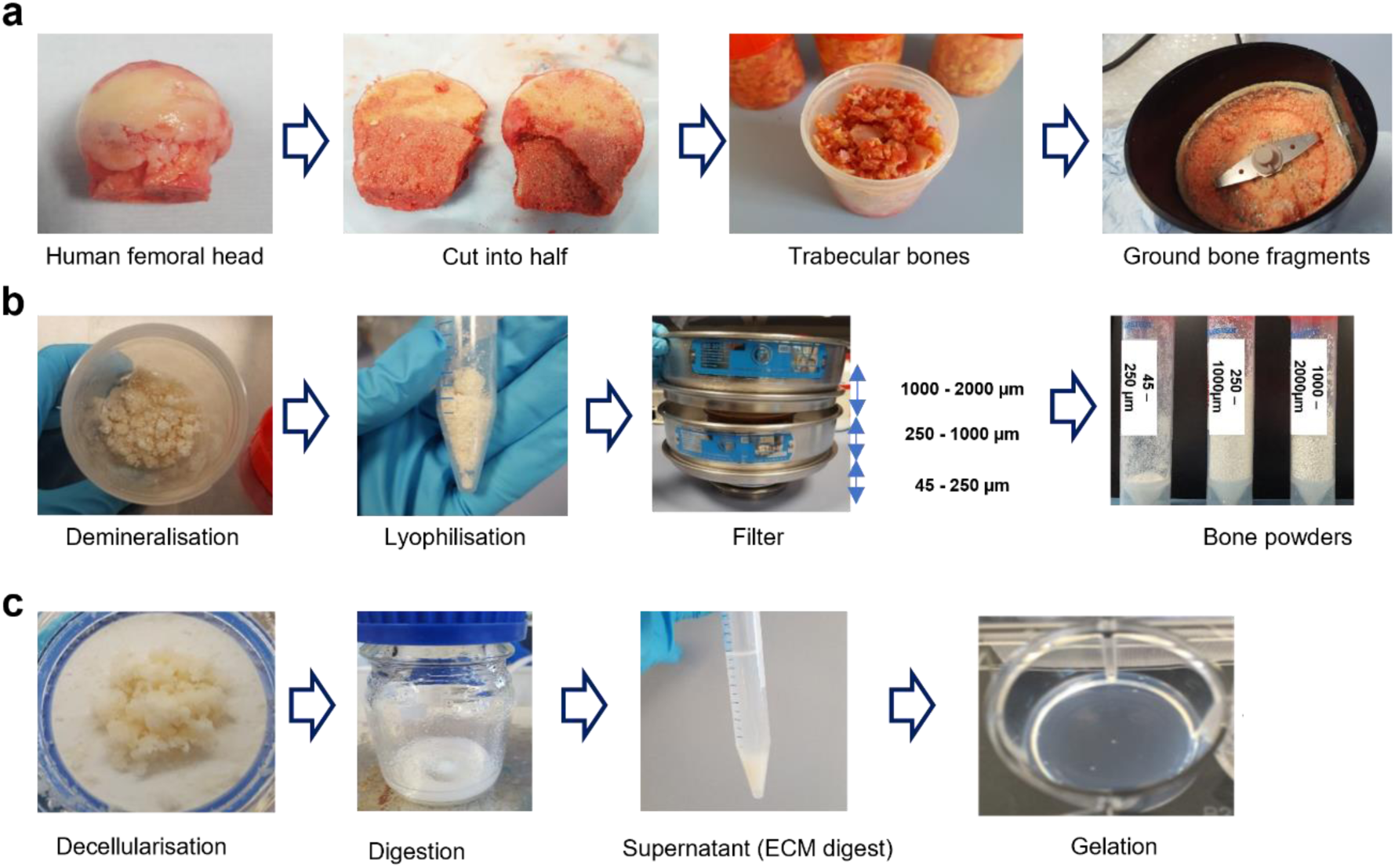
Preparation process of ECM hydrogel derived from demineralised and decellularized human bone. (a) Human femoral heads from female and male patients, aged 65±12 years old were halves, and trabecular bone was collected and fragmented. (b) The fragments underwent demineralisation in 0.5N HCI for 24 hours, followed by lyophilisation. The lyophilised powders were then sieved through stainless steel sieves to obtain powders in the ranges of 45- 250 µm, 250-1000 µm, and 1000-2000 µm. (c) After decellularisation in a Trypsin/ EDTA solution, the powders were digested in a pepsin solution for 3, 5, and 7 days, respectively. The supernatant from the digestion solution was mixed with 0.1N NaOH, 10 × PBS, and 1 × PBS and incubated at 37 °C to induce gelation.

**Figure 2.**
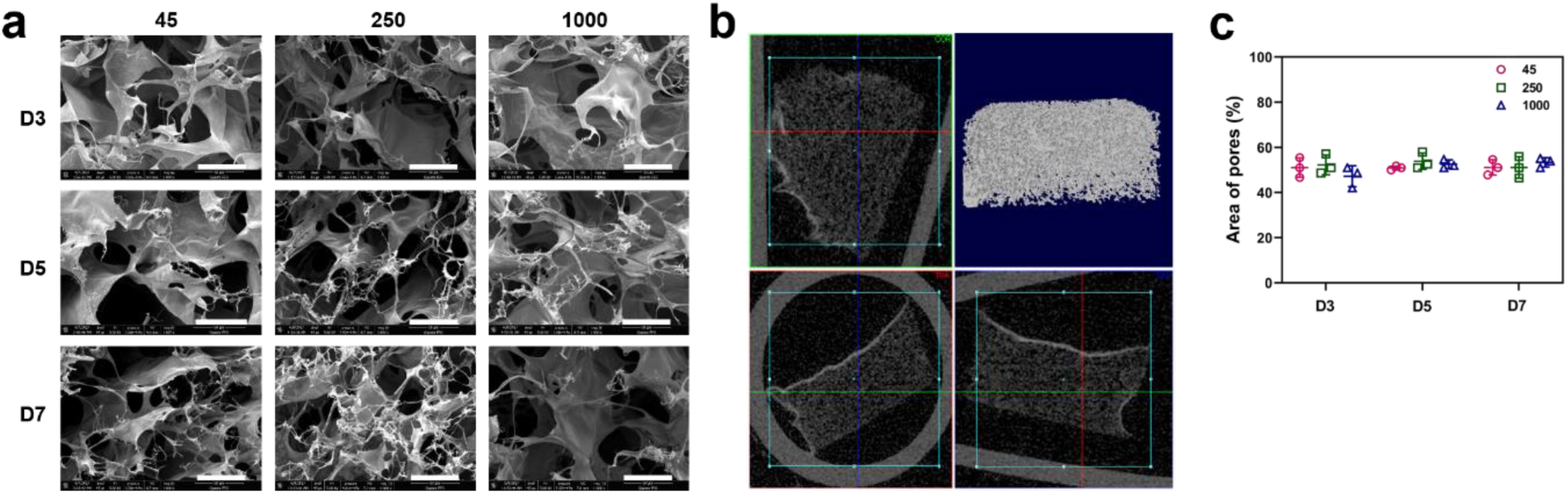
Microstructure analysis of human bone ECM hydrogels using scanning electron microscope (SEM) and micro-computed tomography (μ-CT). (a) SEM images of cross-sections of lyophilised ECM hydrogels were observed by SEM (scale bar = 300 μm), showing the presence of porous and fibrous structures in all types of ECM hydrogels. (b) 3- dimensional visualization of the ECM hydrogel structure using μ-CT demonstrates the porous nature of the hydrogels. (c) Measurement of pore sizes in ECM hydrogels using ImageJ software reveals no significant differences between the hydrogels (n=3). Data represent mean ± SD, N = 4.

**Table 1.**
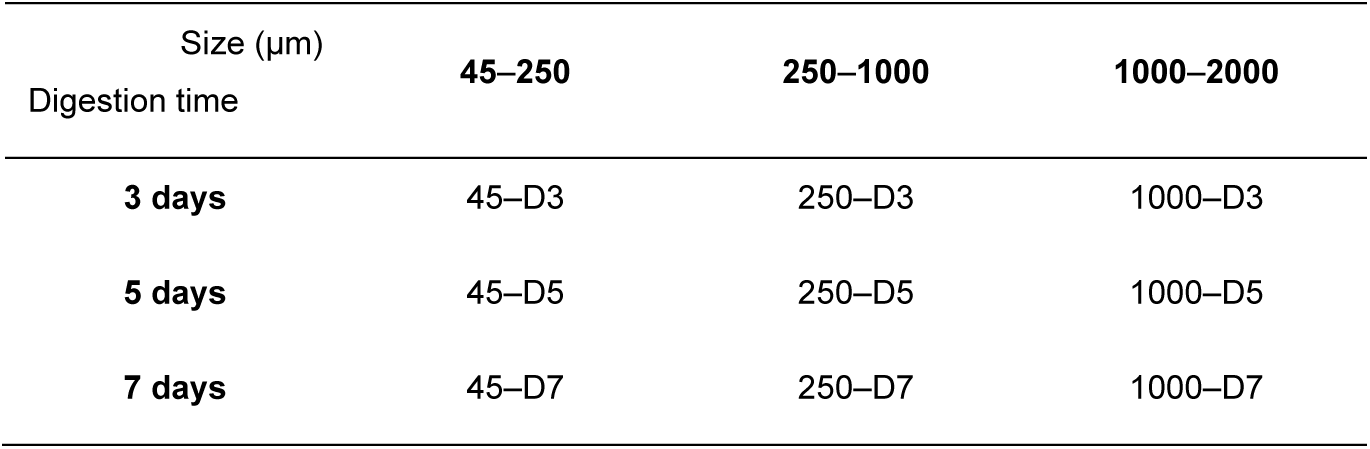
Changes of digestion time and powder size for optimizing bone ECM hydrogels.

| Size (µm) | 45–250 | 250–1000 | 1000–2000 |
| --- | --- | --- | --- |
| Digestion time |  |  |  |
| **3 days** | 45–D3 | 250–D3 | 1000–D3 |
| **5 days** | 45–D5 | 250–D5 | 1000–D5 |
| **7 days** | 45–D7 | 250–D7 | 1000–D7 |

### 2.2 Bone powder size and digestion time affect cellular and protein contents in ECM digests

Different sizes of demineralized bone powders were subjected to digestion with a pepsin solution (ratio of 20 mg powders per unit of pepsin) for durations of 3, 5, or 7 days. Subsequently, the concentrations of DNA and proteins present in the extracellular matrix (ECM) digests were quantified. The concentrations of total proteins, sGAG, and collagen significantly increased as bone powder size decreased and digestion time increased (Figure 3a). Comparing the ECM digest from 1000 – 2000 µm after 3 days of digestion (referred to as 1000–D3, see table 1), it was observed that the samples with the smallest bone powder size and the longest digestion time (45–D7) exhibited a higher solubilization of proteins from the ECM, as indicated by the total protein concentrations (8.5 ± 1.5 mg/ml for 45-D7, 0.6 ±0.16 mg/ml for 1000-D3). The concentration of DNA in 45–D7 ECM digest was approximately 20 ng/ml (∼2.9 ngs of DNA per mg of dried ECM digest), resulting in a content compatible with other types of ECM hydrogels ^[18]^.

**Figure 3.**
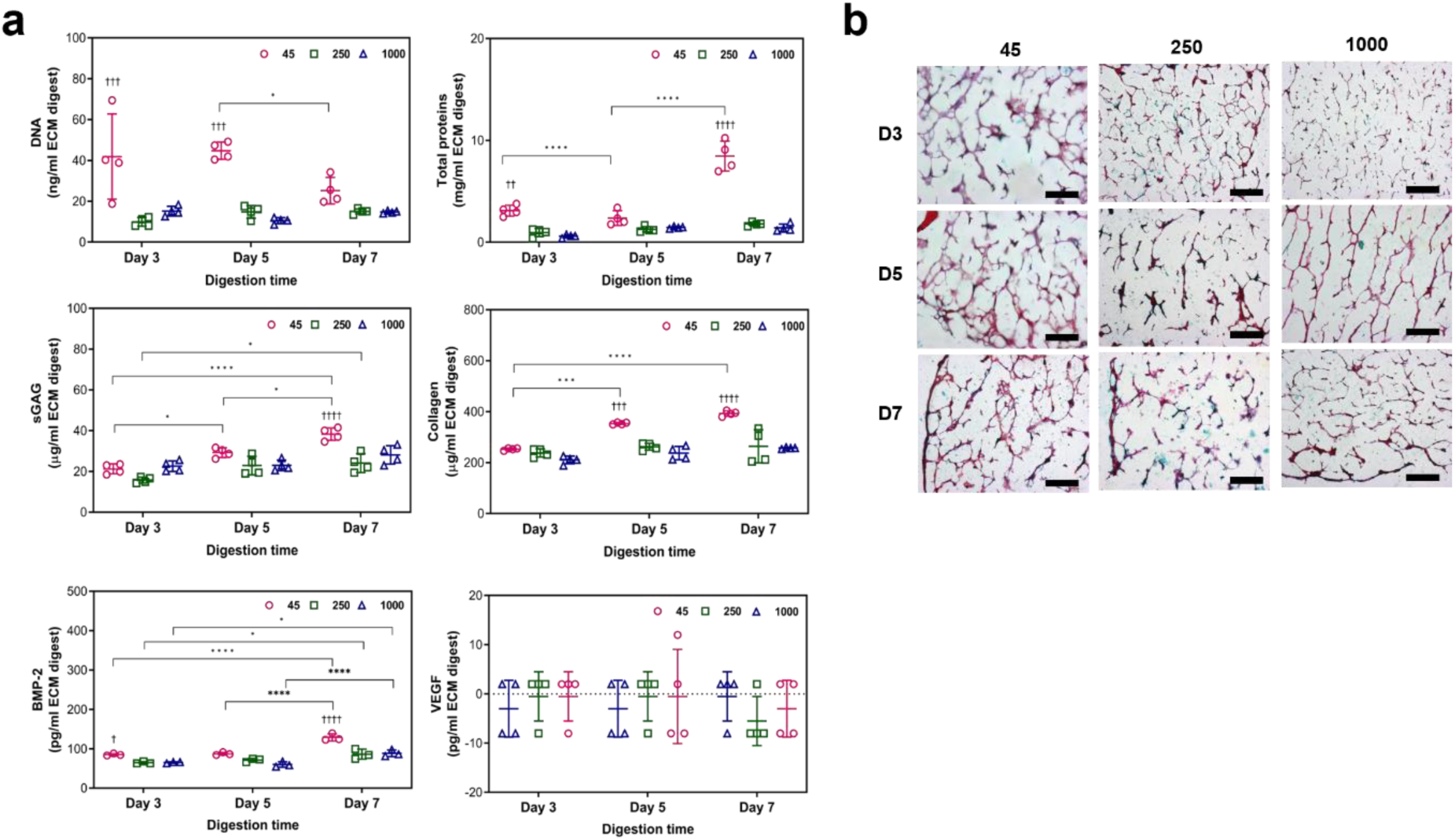
Concentrations of DNA, total proteins, sGAG, collagen, BMP-2 and VEGF in ECM digests and histological staining images of ECM hydrogels. (a) Significantly high concentrations of proteins, including sGAG and BMP-2, were detected in the 45-250 μm powder and 7-day digested ECM digest, compared to other ECM digests (n=4). Statistical significance was determined using a two-way ANOVA test with Tukey’s multiple comparisons test (*p<0.05 **p<0.005 ***p<0.0005****). The † symbol indicates a significant difference within the groups at the same digestion time). Data represent mean ± SD, N = 4. (b) ECM hydrogels were stained with hametoxyline and Alcian blue/Sirius red (scale bar = 100 µm).

To visualize the distribution of proteins within the hydrogels, staining techniques using specific dyes were used. Alcian Blue was used to stain glycosaminoglycans (blue coloration). Sirius Red for collagen (pink/red coloration) and hematoxylin were used to stain nuclei (black coloration). The hydrogels exhibited prominent staining for collagen and GAGs (Figure 3b), while nuclei staining was less pronounced. ECM hydrogels derived from small bone powder sizes and treated for longer digestion times exhibited stronger staining of collagen and GAGs across larger areas. However, the nuclei staining was barely visible, presenting as a black coloration. These results were consistent with the cellular and protein contents observed in the hydrogels (Figures 3a). Furthermore, as depicted in Figure 2, the staining images revealed the presence of porous structures in all the hydrogels.

BMP-2 molecules were detected in all of the ECM hydrogels with the 45-D7 hydrogel exhibiting the highest concentration of BMP-2 (129 pg/ml BMP-2 ± 9.1, Figure 3a). However, no VEGF was detected, possibly due to the fact that VEGF is produced by hypertrophic chondrocytes, while BMP-2 is primarily secreted into the bone matrix by pre-existing osteoblasts, osteocytes, and endothelial cells ^[19, 20]^. The presence of BMP-2 in the ECM hydrogels suggests their significant potential for use in bone repair applications.

### 2.3 Bone powder size and digestion time affect gelation speed, rheological strength, and degradation of ECM hydrogels

To investigate the influence of bone powder size and digestion time, on the gelation and rheological properties of ECM hydrogels, turbidimetric gelation kinetics analysis was undertaken. The gelation kinetics of all the human bone ECM hydrogels exhibited a sigmoidal pattern (Figure 4a). Hydrogels derived from bone powder size ranging from 45 to 250 µm (45-D3/D5/D7), regardless of digestion time, demonstrated significantly sharper slopes and shorter t_lag_ and t_1/2_ compared to those derived from 250 – 1000 µm and 1000 – 2000 µm (Figure 4c) bone powders (Figure 4c). Specifically, the 45–D3/D5/D7 hydrogels displayed gelation approximately 7 ± 3 min after incubation at 37 °C, while the 250–D3/D5/D7 and 1000–D3/D5/D7 hydrogels exhibited gelatin times of 16 ± 0.4 and 20 ± 0.5 min, respectively. For the gelation process, various ECM digests were mixed with the ECM gelation buffer, and the resulting pre-gel ECM solutions were placed in individual tubes and gently inverted. Some pre-gel solutions for 250–D3, 250–D5, 1000–D3 and 1000–D5 were immediately flowed down, while other samples remained stable in the tubes (Figure 4b). After 1 hour of incubation at 37 °C, with the exception of 250–D3 and 1000–D3 samples, all the ECM hydrogels maintained their structure in the inverted tubes.

**Figure 4.**
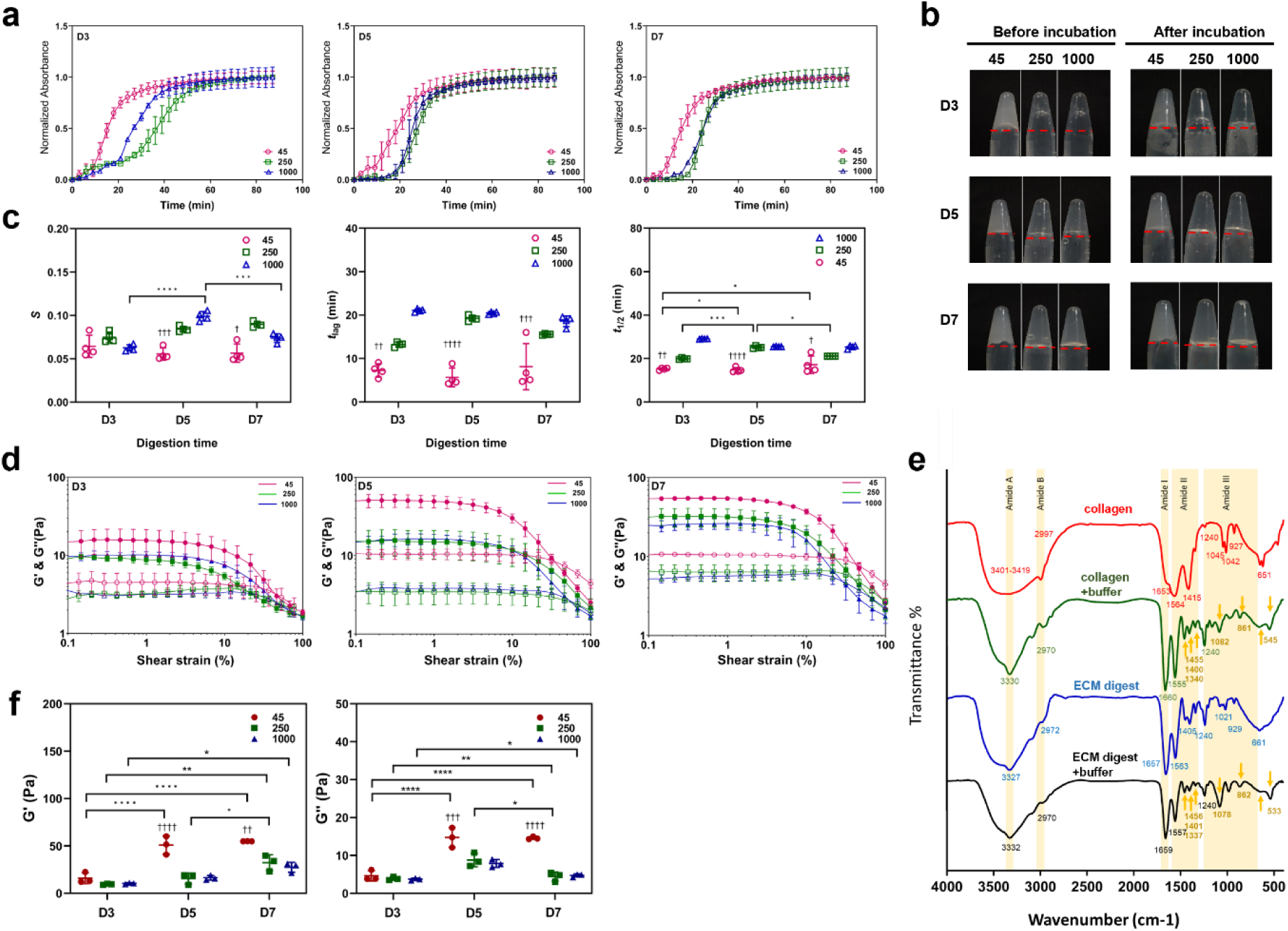
Gelation and rheological characterizations of ECM hydrogels from various powder sizes and digestion time. (a) Turbidimetric gelation kinetics of ECM hydrogels. pH-neutralized ECM pepsin digests were added to the wells of a pre-warmed 96-well plate (37 °C), and the absorbance at 405 nm was measured at 3-minute intervals (n=4). The values were normalized between 0 (the initial absorbance) and 1 (the maximum absorbance). (b) Photograph images of each gel after pH neutralization in inverted tubes before and after incubation at 37 °C. The red line indicates the initial volume of the gel before inversion. (c) S (speed of gelation), tlag (time to start gelation), and t1/2 (time to reach 50% maximum turbidity) were calculated based on turbidimetric curves. (d and f) Storage modulus (G’) and loss modulus (G’’) were monitored as hydrogels underwent an amplitude sweep of 0.1-100% strain at a constant angular frequency. Data represent means ± standard deviation for n=3. (e) Fourier transform infrared spectroscopy (FTIR) spectra of ECM digest and collagen with and without ECM buffer were analyzed. The spectrometer’s detection range was 399-4000 cm-1, and the data were measured with an interval of 0.96 cm-1 at room temperature. Statistical significance was determined using a two-way ANOVA test with Tukey’s multiple comparisons test (*p<0.05 **p<0.005 ***p<0.0005****). The † symbol indicates a significant difference within the groups at the same digestion time). Data represent mean ± SD, N = 4.

Furthermore, we evaluated the rheological characteristics of the hydrogels to determine the impact of bone powder size and digestion duration on the strength of human the bone ECM hydrogels (Figure 3d and f). The storage (G’) moduli of the hydrogels derived from 45 – 250 µm bone powders, regardless of digestion time, were significantly higher compared to those derived from 250 – 1000 µm and 1000 – 2000 µm bone powders. The highest storage modulus was observed in the 45-D7 hydrogels (55.0 ± 0.1 Pa), which was three-times higher than the modulus of 45-D3 (16.90 ± 5.7 Pa). No significant difference was observed between the modulus of 45-D7 and 45-D5 (50.9 ± 9.6 Pa) hydrogels. In case of ECM hydrogels derived from 250 – 1000 µm and 1000 – 2000 µm bone powders, there were no clear difference in storage and loss moduli was observed between D3 and D5 hydrogels. However, the moduli of 250-D5 were significantly higher than those of 1000-D5. These rheology results suggest that a digestion time of 5 days enabled solubilization of a variety of proteins form the ECM to generate stiff hydrogels using 45 – 250 µm bone powders. However, more time was required to digest proteins from large bone powders.

To investigate the impact of ECM buffer on ECM gelation, we evaluated the chemical structures of type I collagen and ECM digest with and without ECM buffer using Fourier Transform Infrared Spectroscopy (FTIR) (Figure 4e). The spectra of all samples exhibited distinct patterns across six regions; 3400-3420 cm^-1^ (amide A), 3000-2970 cm-1 (amide B), 1660-1657 cm^-1^ (amide I), 1557-1563 cm^-1^ (amide II), and 1240-670 cm^-1^ (amide III), which are consistent with previously reported patterns in the literatures ^[21, 22]^.

The broad band observed at 3327-3419 cm^-1^ corresponds to the stretch –OH groups in collagen. Stretch vibration of C–H were observed at 2970-2997 cm^-1^. The absorption band at 1650 cm^-1^ in Amide I region indicates that stretch vibration of C=O in the polypeptide backbone of the protein. The peaks around 1550 cm^-1^ represent the amide II bands corresponding to N–H bending. The characteristic peaks in the amide A, B and I regions were present in all samples, regardless of the addition of ECM buffer. However, the effect of ECM buffer on collagen and ECM digest was evident in the peaks in the amide II and III regions. Lower intensity absorption bands were observed in collagen and ECM hydrogels (with ECM buffer) compared to collagen and ECM digest at around 1450, 1400, 1340, and 660 cm^-1^. These bands were attributed to CH2 bending, CH2 wagging of proline, COO- symmetrical stretch, and O=C–N bending, respectively, indicating a greater disorder associated with the loss of the triple helix state of collagen ^[23]^. Peak intensities of around 1080 cm^-1^and 860 cm^-1^, indicative of ether linkage (C–O–C), were increased in the collagen and ECM gels treated with ECM buffer. This observation suggests the presence of NaCl and NaOH in the buffer plays an important role in the gelation of collagen and ECM hydrogels by forming a linkage between the polypeptide chains of collagen.

To validate protein efficacy on hydrogel stability in vitro, we evaluated the amounts of released proteins from the ECM hydrogels after incubating them in PBS for 24 hours, followed by collagenase-PBS for an additional 24 hours at 37 °C (Supplementary Figure S2). Following 1 hour of incubation in PBS, approximately 20-25 % of proteins were released from the ECM hydrogels, and this percentage remained consistent even after 23 hours. However, upon addition of the collagenase-PBS solution, the protein amounts from all the hydrogels rapidly increased following enzymatic stimulation by collagenase. Furthermore, it was noted the ECM hydrogels derived from smaller bone powder samples exhibited a significantly slower release pattern (Supplementary Figure S2a), indicating that these hydrogels undergo degradation at a slower rate compared to the larger bone powder-derived hydrogels. Among the 45 bone powder-sized ECM hydrogels, prolonged digestion times resulted in slower degradation, but this trend was not observed for the 250 and 1000 bone powder samples. Since their protein concentrations were comparable, (Figure 3a), it appears protein concentration affects the degradation of ECM hydrogels. These results suggest bone powder size and digestion times play a crucial role in determining the gelation and rheological properties of ECM hydrogels, as well as their degradation capacity.

### 2.4. Proteomic analysis of human bone ECM digests

We have demonstrated that ECM hydrogels derived from smaller bone size and treated for prolonged digestion times enhance a higher concentration of proteins from demineralised and decellularised human bone ECM, as indicated by total protein, collagen and sGAG assays (Figure 3a). However, accurately capturing the complex compositions of the ECM remains a challenge. To validate the effect of bone powder size and digestion time on the quality and quantity of ECM, an intensive proteomic analysis of the ECM hydrogels was undertaken. Following ECM digestion, a maximum of 246 proteins were identified from the ECM digests, including 52 core-matrisome proteins (collagens, glycoproteins, and proteoglycans) and 31 matrisome-associated proteins (ECM regulators, secreted factors, and ECM affiliated proteins), using the ECM-specific categorization database, MatrisomDB 2.0 ^[24, 25]^. The percentage of categorized proteins was calculated based on Score sequest HT, which scores and sums each peptide for a protein.

The impact of bone powder size and digestion time was crucial in determining the proportion of categorized proteins in ECM digests, as demonstrated in Figure 5 a and b. While the protein quantification assay results (Figure 3a) clearly show an increase in protein concentration with decreasing bone size and increasing digestion time, this trend was not consistently observed in the proteomic results. Interestingly, the percentages of all the categorized proteins from ECM digests treated for 5 days were dramatically higher or lower than those treated for 3 days and 7 days, regardless of bone powder size (Figure 5b). Specifically, the ECM digest derived from 45 – 250 µm bone powder size and treated for 5 days (45–D5) showed a lower percentage of collagens but higher percentages of other matrix proteins. Figure 5c illustrates the percentages of categorized proteins in ECM digests from the different sizes of bone powders after 5 days of incubation (45–D5, 250–D5, and 1000–D5). As bone powder size increased, an observed increase in proportion of core-matrisome proteins was noted and a corresponding decrease in non-matrisome proteins. Thus, within the core-matrisome category, 45–D5 displayed a much lower percentage of collagens (45–D5; 69.5%, 1000–D5; 91.3%), but a higher percentage of glycoproteins and proteoglycans were detected (Figure 5c). Furthermore, a range of proteins categorized as core-matrisome, matrisome-associated, and non-matrisome were identified in 45–D5 ECM digest compared to ECM digests derived from higher bone powder sizes (250–D5 and 1000–D5, Figure 3d and e). Furthermore, there were more proteins exclusively present in 45–D5 ECM digest (Figure 5d and e (iii)), including well-known bone formation-related molecules such as osteoglycin, osteocalcin, insulin-like growth factor, and transforming growth factor. These findings highlight that smaller bone powders result in the digestion of a higher concentration of proteins with a wider range of protein types, offering significant promise for bone regeneration.

**Figure 5.**
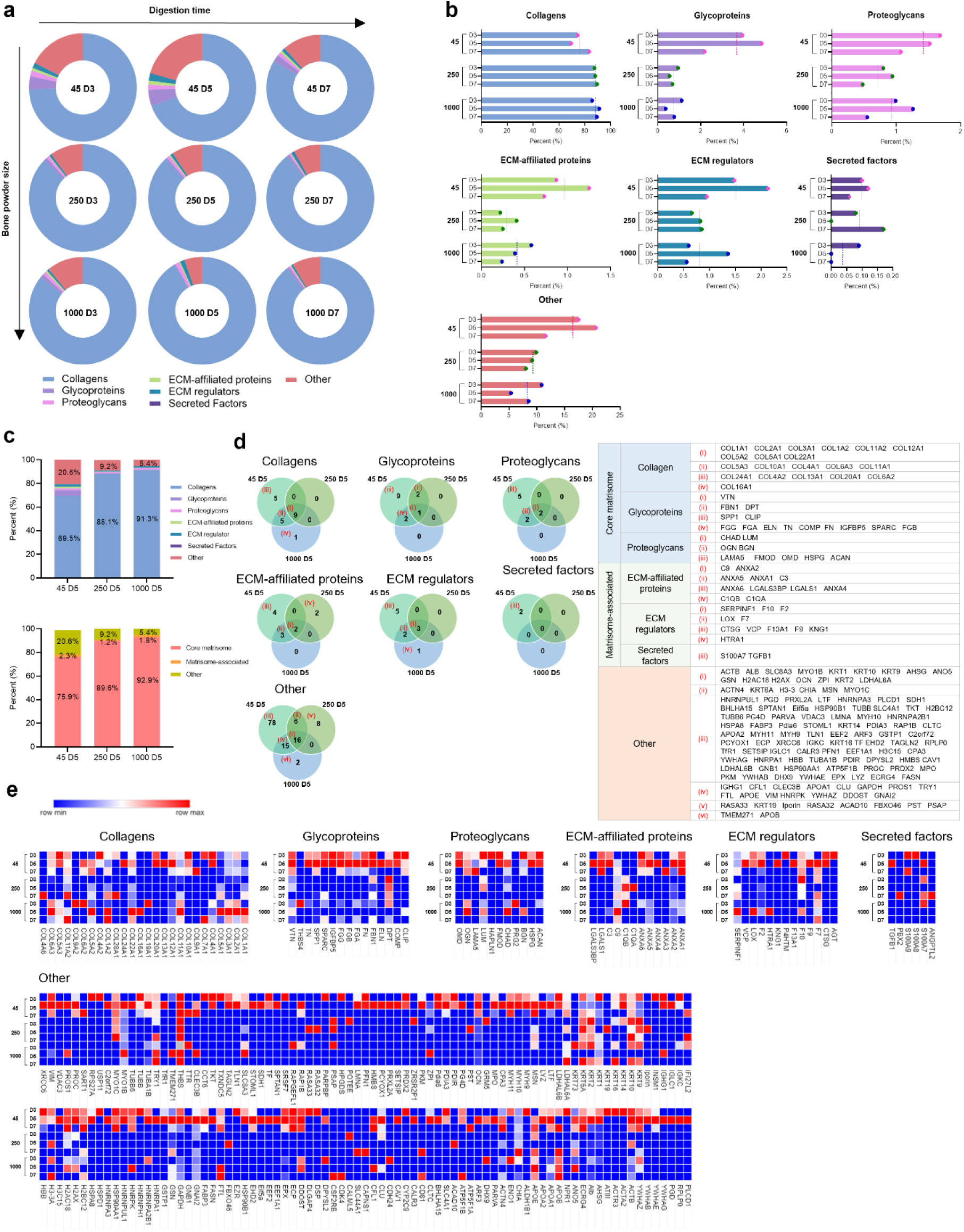
Comparison of the total matrisome subcategories of proteins in human bone ECM digests. (a and b) illustrate the percentages of proteins categorized under different matrisome subcategories in ECM digests. The dashed lines in (b) represent average percentages from the group. ECM digests derived from various sizes of bone powders after 5 days of digestion exhibit a significant impact on the protein proportions compared to those from shorter or longer incubation times. (c) presents the percentages of categorized proteins from 45-D5, 250-D5 and 1000-D5. (d) shows a Venn diagram depicting the percentage of matrisome subcategories of proteins in various sizes of bone powders after 5 days of digestion. Smaller bone powder exhibits a lower presence of collagens but a higher presence of other protein types. The details and proportions of the proteins are listed in the table (d) and heatmap (e).

### 2.5 Human bone ECM hydrogels improve bone marrow-derived stromal cell function

We investigated the ability of human bone ECM to support human marrow-derived stromal cells (HBMSCs) due to their biological potential in modulating cell response and promoting bone tissue regeneration. Initial studies focused on evaluating HBMSCs functions, including attachment, spreading, migration viability and proliferation, when cultured in human bone ECM hydrogels derived from various human bone powder sizes and treated for 7 days (45-D7, 250-D7 and 1000-D7). Regardless of the bone powder size examined, cells exhibited excellent bone particle attachment and were observed to spread extensively within 30 minutes of incubation, evidenced by the organization of stress fibers and the formation of filopodia associated with the gel surfaces (Figure 6a). To compare cell behaviors with a control, a droplet of ECM hydrogels was placed in the centre of a tissue culture plate (TCP), and time-lapse recordings performed. (Figure 6b and Supplementary Video 1). Cells cultured on TCP and the ECM hydrogels exhibited excellent cell spreading and intercellular interactions. Interestingly, some cells on the TCP were observed to move towards the ECM hydrogels, while cells on the ECM hydrogels did not migrate towards the TCP.

**Figure 6.**
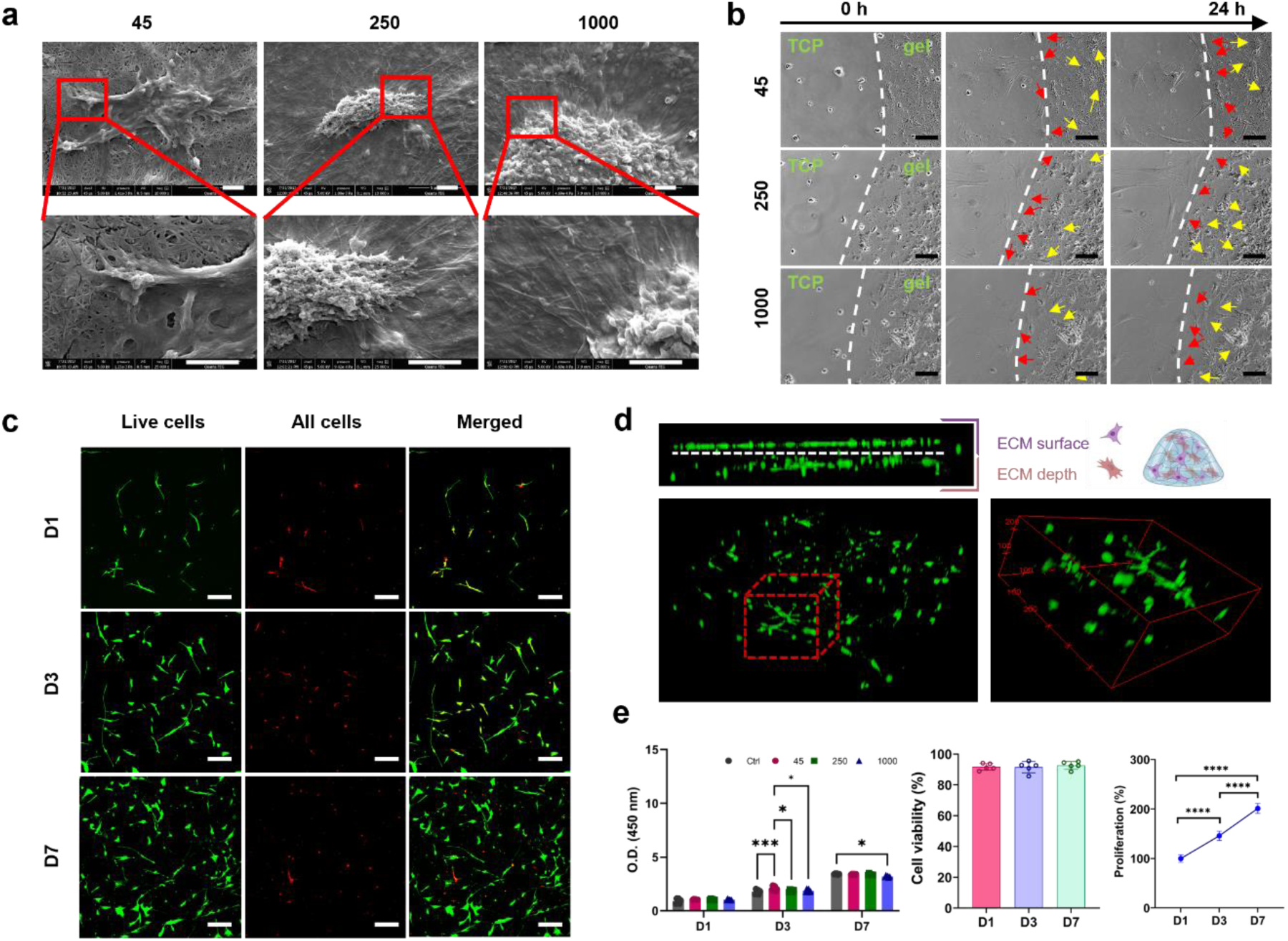
Behaviour of human bone marrow-derived stromal stem cells (HBMSCs) on human bone ECM hydrogels. (a) HBMSCs were observed to be well attached and spread on the ECM hydrogels, irrespective of the hydrogel source from various bone powder sizes (scale bar = 2 µm). (b) ECM hydrogels were placed in the center of tissue culture plate (TCP), and after seeding HBMSCs, their behaviours were observed using a time-lapse microscope. Some cells from the TCP migrated towards the ECM hydrogels (indicated by red arrows), while cells on the ECM hydrogels spread on top of the hydrogels (indicated by yellow arrows) (scale bar = 50 µm). (c and d) Cells stained with Calcein AM and DiD (green and red, respectively) were cultured in ECM hydrogels for 1, 3, and 7 days. (e) Cell viability and proliferation were evaluated using confocal imaging analysis and the WST-1 assay (n=5). The cell population almost doubled within 7 days, and there was no significant difference in ECM hydrogels derived from various bone powder sizes. Statistical significance was determined using a one-way test with Tukey’s multiple comparisons test (*p<0.05 **p<0.005 ***p<0.0005****).

The presence of a wide variety of proteins in the ECM hydrogels (Figure 5) may have promoted cell migration/chemotaxis towards the hydrogels. However, despite significant differences in the release patterns of proteins from the ECM hydrogels (Supplementary Figure S2), no significant differences in cell migration behavior were observed across the different ECM hydrogels. These results suggest that linear boundary of ECM hydrogels stimulated cells to migrate together with an elongated morphology ^[26, 27]^. For cells initially seeded on the ECM hydrogels, the porous and fibrous structures of the hydrogels provided an attractive environment for cell exploration (Figure 2 and Supplementary Figure S1) ^[28]^. Cell morphology on the ECM hydrogels were distinctly different compared to cells on TCP and ECM digest-coated TCP as observed following staining for alkaline phosphatase expression (Figure 7a). Cells were observed to invade into the porous and fibrous microenvironment of the ECM hydrogels, rather than remaining within the surface. Furthermore, these ECM hydrogel structures promoted cell proliferation, with a doubling in cell numbers within 7 days and excellent viability (Figure 6c, d and e). However, we didn’t observe any significant effect of varying bone powder sizes of ECM hydrogels on HBMSCs proliferation.

**Figure 7.**
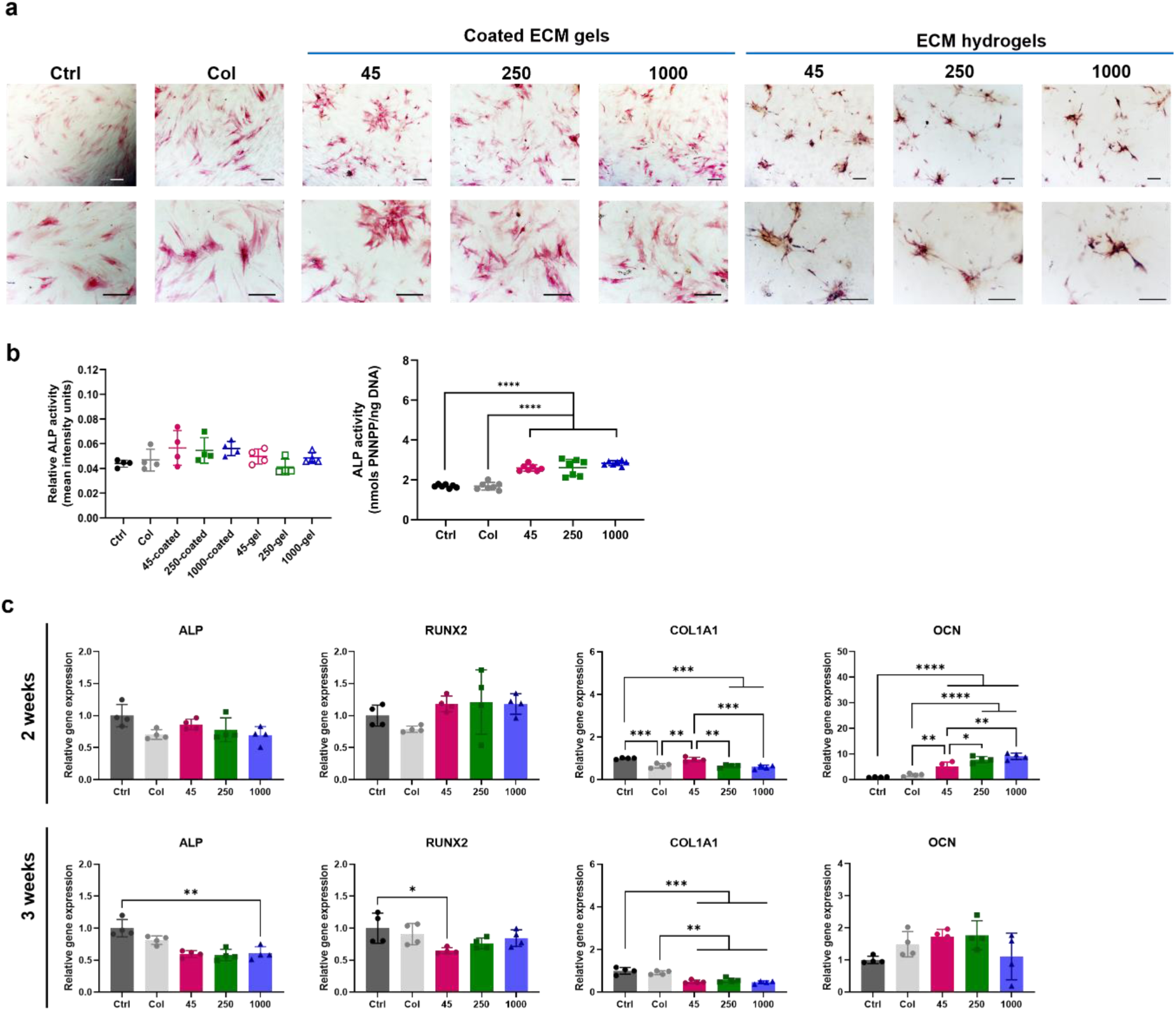
Impact of human bone ECM hydrogels on HBMSCs osteogenic differentiation. (a) HBMSCs morphology and alkaline phosphatase (ALP) staining density/color differed notably between the 2D and 3D settings. (b) ECM hydrogels derived from bone powder sizes of 45–250 µm remarkably enhanced the early and late responses of HBMSCs differentiation compared to collagen gels and larger bone powder-sized ECM hydrogels. Statistical significance was determined using a one-way test with Tukey’s multiple comparisons test (*p<0.05 **p<0.005 ***p<0.0005****). Data represent mean ± SD, N = 4.

Osteogenic differentiation of HBMSCs was assessed on ECM digest-coated surfaces (2D) and ECM hydrogels (3D) using ALP staining and ALP/DNA assays after 1 week of HBMSCs culture (Figure 7a and b). Compared to collagen type 1 gel (Col) and control (Ctrl, TCP), ALP staining activities on ECM digest coated surfaces, regardless of ECM digests from different powder sizes, showed a slight increase compared to the control and collagen-coated surfaces. Interestingly, the cell morphology and staining density/color differed notably between the 2D and 3D settings. For instance, cells on the 2D surfaces exhibited widespread pink staining, while on the 3D hydrogels, cells formed aggregates with dark pink/purple and brown coloration. As the staining density for ALP activity assessment was based on the pink color, no significant differences were observed. However, when ALP activity (nmols PNNPP/ng DNA) was assessed using ALP assay and normalized by DNA content, all the ECM hydrogels significantly enhanced ALP activity compared to the control and collagen-coated surface materials.

To validate the osteogenic potential of human bone ECM hydrogels, the expression levels of early markers (ALP, RUNX2, and COL1A1) and a late (OCN) marker of osteogenic differentiation were evaluated after culturing HBMSCs on ECM hydrogels for 2 and 3 weeks (Figure 7b). After 2 weeks, the ECM hydrogels showed a significant increase in RUNX2 expression compared to collagen gel and control (TCP). Furthermore, the expression level of COL1A1 in ECM hydrogels derived from 45 – 250 µm sized bone powder was significantly enhanced than observed in collagen gel and ECM hydrogels from larger bone powder samples. Among all the early markers, it was observed that the expression levels from the 45 ECM hydrogels were found to be significantly higher than those from the collagen gel after 2 weeks, but expression levels from the 45ECM hydrogel were lower than the collagen gel after 3 weeks. Several studies have shown that ALP, RUNX and COL1A1 activity increase during the first two weeks of osteogenic differentiation and subsequently decline to facilitate mineralization ^[29-31]^. These results suggest the 45 ECM hydrogel has the potential to modulate the early differentiation response of HBMSCs.

After 2 weeks, all the ECM hydrogels exhibited significant upregulation of the late marker OCN gene compared to the control and collagen gel. However, after 3 weeks, there was a notable decrease in OCN expression compared to expression after 2weeks, although the levels from the 45 and 250 ECM hydrogels remained higher than those from the control and collagen gel. Interestingly, the proportion of collagens in the ECM hydrogels, particularly those derived from lower bone powder sizes, was much lower and consisted of various core-matrisome, matrisome-associated, and non-categorized proteins (Supplementary Figure S3). Collagen Type I has a key role in promoting osteogenic differentiation ^[32, 33]^. This suggests that the presence of various proteins in the ECM is crucial for enhancing osteogenic differentiation and the importance of human bone powder size in generating hydrogels consisting of a variety of ECM proteins.

## 3. Conclusion

Decellularized ECM hydrogels derived from human bone tissue have garnered significant interest in the field of tissue engineering given their native components and structures. This study represents the first successful production of a hydrogel from demineralized and decellularized human bone extracellular matrix. Moreover, we have demonstrated the critical role of bone powder size and digestion duration in the generation of human bone ECM hydrogels and the subsequent modulation of gelation and rheological properties, as well as the quantity and composition of digested proteins within the bone ECM. Notably, smaller bone powder sizes facilitate the digestion of a wider range of ECM proteins offering significant potential as biologically active relevant hydrogels for bone tissue engineering applications.

## 4. Experimental Section

### Demineralization of human trabecular bone

Human femoral heads were collected from haematologically normal patients undergoing elective hip arthroplasty. Only tissue samples that would have been discarded were used following informed consent from the patients in accordance with the Southampton & South West Hampshire Local Research Ethics Committee (Ref: 194/99/w). The femoral heads from female and male patents, aged 65±12 years old were cut using an isomet cutting machine (Buehler LTD. US), and trabecular bones from the femoral heads were collected using a bone nipper (Figure 1a). The collected trabecular bones were frozen in liquid nitrogen and ground in a commercial coffee grinder. To remove residual fats from the bones, the ground bones in 50 ml tubes were washed with phosphate-buffered saline (PBS, Lonza, Slough, UK) containing 2% penicillin/streptomycin (P/S, Sigma–Aldrich, Poole, UK) until the tubes were no longer smeared with fat.

Ground human trabecular bones were demineralised using a modification of previously reported methods ^[15]^ (Figure 1b). The bone fragments were treated in a 0.5 N HCl (Fisher Scientific) at room temperature for 24 hours. To solubilize the lipids in the bone, the fragments were added to a 1:1 mixture of chloroform and methanol for 1 hour, followed by rinsing in methanol and then deionized water several times. After lyophilization, the demineralized bone matrix (DBM) fragments were ground using a homogenizer (T10 basic Ultra-turrax, IKA LTD, Oxford, UK). The powders went passed through stainless steel sieves with 45, 250, 1000, and 2000 µm pore size (Fisher Scientific) to obtain the DBM powders with 45 – 250 µm, 250 – 1000 µm, and 1000 – 2000 µm sizes.

### Decellularization and digestion

The DBM powders were decellularized in a solution of 0.05% trypsin with 0.02% ethylenediaminetetraacetic acid (trypsin/EDTA, Lonza) at 37 °C and 5% CO_2_ incubator under agitation for 24 hours (Figure 1c). The powders were collected using cell strainer (40 µm, Fisher Scientific) and rinsed in PBS twice and in deionized water twice, followed by overnight freeze-drying. Pepsin (Sigma–Aldrich) was dissolved in 0.01N HCl (1 mg/ml). The demineralized and decellularized bone powders with various sizes (45- 250 µm, 250-1000 µm, 1000-2000 µm) were separately added to the pepsin solution and enzymatically digested under agitation for 3, 5, and 7 days, respectively, at room temperature (Table 1). The final concentration of the bone powders in the pepsin solution was 20 mg/ml, and the solutions were then centrifuged at 2,000 rpm at 15 minutes to obtain supernatants referred to as ECM digests. The supernatants were collected and stored at -20 °C until required.

### Gelation of human bone ECM hydrogel

The ECM hydrogels were induced by neutralizing the pH and salt concentration of the ECM digests, followed by incubation at 37 °C, as previously described ^[15, 34]^. Briefly, ECM digests were mixed with one-tenth of the digest volume of 0.1 N NaOH, one-ninth of the digest volume of 10× PBS, and then diluted with 1× PBS (referred to as ECM gelation buffer) on ice to create a 16 mg/ml ECM gel. Subsequently, the neutralized pre-gel solution was incubated for 1 hour at 37 °C.

### Microstructure observation

Microstructure of human bone ECM hydrogels were examined using scanning electron microscopy (SEM). The pre-gel ECM solutions were neutralized and incubated at 37°C for 1 hour, followed by freeze drying to dehydrate the gels. The dried gels were coated with platinum and analyzed using an SEM (FEI Quanta 200) at accelerating voltages of 5 kV. Images were captured at magnifications of 55 ×, 200 ×, 1000 × and 4,000 ×. The area of pores was measured using ImageJ software (n=3).

Additionally, the freeze-dried ECM hydrogels contained in microcentrifuge tubes were individually scanned using Micro-Computed Tomography (µ-CT, SkyScan 1176 (Bruker, Kontich, Belgium). The scanning parameters were set to an X-ray source voltage of 50 kV, current of 500 μA, exposure time of 496 ms, and a voxel size of 18 μm.

To observe the structure of the hydrogels and the presence of various proteins derived from human bone, histological staining was performed. The ECM hydrogels were embedded in optimal cutting temperature compound (OCT, Fisher Healthcare™ Tissue-Plus™) and frozen at -20 °C. Sections in thickness of 8 µm were obtained using a Cryostat (CM 1850, Leica Biosystems) and mounted with super cryo-mounting medium type R3 (SCMM R3, Section lab Co.Ltd, Japan). The sections were then stained using Alcian Blue/Sirius Red staining, including haematoxylin. Images were obtained using a microscopy (Axiovert 200, Zeiss).

### Quantification of cellular, protein, growth factor contents

The cellular content of the various ECM digests was determined using a Quant-iTTM PicoGreen dsDNA assay kit (Invitrogen, Fisher Scientific, Paisley, UK). Following the manufacturers’ protocol, 100 µl of ECM digest samples and DNA standards ranging from 0 to 1000 ng/ml were added to each well of a 96 well plate and mixed with 100 µl of the working solution (n = 4). The plate was covered with foil and incubated at room temperature for 5 minutes. The sample fluorescence was measured at excitation and emission wavelengths of 475 nm and 550 nm, respectively, using a microplate reader (Glomax, Promega, Southampton, UK). Based on a standard curve, the concentration of DNA in the ECM digests was determined.

The total protein concentration of the ECM digests was measured using a DC protein assay kit (Bio-Rad, Watford, UK) following the manufacturer’s instruction (n = 4). Briefly, 5 µl of protein (bovine serum albumin) standard and the digest samples were added to each well followed by addition of 225 µl of the working reagent. After 15 minutes, the absorbance was read at 750 nm using the microplate reader.

The collagen and sulphated glycosaminoglycan (sGAG) concentrations of the digests were also quantified using the Sircol collagen assay and BlyscanTM Sulfated Glycosaminoglycan assay (Biocolor Lt, Carrickfergus, UK). The digests were diluted with a pepsin solution and analyzed for the collagen and sGAG concentrations (n = 4) according to the manufacturer’s protocol.

The quantification of bone morphogenetic growth factor (BMP)-2 and vascular endothelial growth factor (VEGF) content was conducted using human BMP-2 and VEGF Quantikine enzyme-linked immunosorbent assay (ELISA) Kits (R&D, Bio-Techne, Abingdon, UK), respectively. Those assays were performed according to the manufacturer’s instruction (n=4) and the concentration of growth factors in the ECM digests was determined based on their standard curves.

### Gelation characterization – Tube inversion and turbidimetric studies

The gelation property of human bone ECM hydrogels was determined through tube inversion and turbidimetric gelation assays. For the tube inversion study, the neutralized pre-gel solutions (16 mg/ml) obtained from various bone ECM powder sizes and digestion times were placed in 1.5 ml of Eppendorf tubes and the volume of each solution in the tubes was marked. After inverting the tubes, images were taken using a digital camera. Subsequently, the tubes were incubated at 37 °C for 1 hour, inverted again, and images were taken.

Next, turbidimetric gelation kinetics of the ECM hydrogels were evaluated spectrophotometrically as previously described ^[34, 35]^. The neutralized pre-gel solutions were prepared on ice, and 100 µl of the solution was transferred to a pre-cooled 96 well-plate. The plates were placed in a pre-warmed microplate reader at 37°C, and the turbidity of each well was measured at a wavelength of 450nm every 2 minutes for an hour (n = 4). The absorbance values for each well were recorded and averaged.

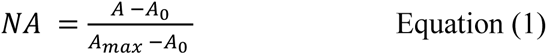

These readings were then scaled from 0 (at time 0) to 100% (at maximum absorbance) for normalizing absorbance (NA), as shown in Equation (1), where A is the absorbance at a given time, A0 is the initial absorbance, and Amax is the maximum absorbance. The time needed to reach 50% of the maximum turbidity measurement (e.g. maximum absorbance value) was defined as t1/2 and the lag time (tlag) was calculated by the intercept of the linear region of the gelation curve with 0% absorbance. The speed (S) of the gelation based on turbidimetric measurements was determined by calculating the maximum slope of the growth portion of the curve as previously described ^[34]^.

### Gelation characterization – Rheological characterization

The rheological characteristics of human bone ECM hydrogels were determined using a rheometer (MCR 92, Anton Paar, UK). 1 ml of pre-gel solution was loaded onto a pre-warmed (37 °C) 25 mm parallel plate and incubated at 37 °C for 1 hour under humanify condition. An amplitude sweep experiment was conducted for the human bone ECM at 0.01-200% stain at a constant angular frequency of 1 rad/s.

### Gelation characterization – Fourier transform infrared spectroscopy (FTIR) analysis

The neutralized pre-gel ECM and collagen solutions (with ECM buffer), ECM digest, and collagen (without ECM buffer) were analyzed by FTIR (FT-IR-4700, Jasco, Maryland, USA). 16 mg/ml of ECM digest and ECM pre-gel solution and ECM digest, and 300 µg/ml type I collagen and collagen solution (buffer) were drop casted and dried overnight. The detection range of the spectrometer was 399 – 4000 cm^-1^, and the data were measured with an interval of 0.96 cm-1 at room temperature. The raw FT-IR data was processed with a spectrum software (Spectra Manager™ Suite, Jasco) for smoothing and baseline correction.

### Proteomic analysis – sample preparation

ECM digests derived from various bone powder sizes and treated digestion times were prepared for mass spectrometry. Briefly, ECM digests were solubilized in 5 M urea with 50 mM of (NH_4_)HCO_3_ and vortex for 30 s. To reduce disulfide bonds, 5 mM dithiothreitol was added to the samples, followed by 1 hour incubation under agitation (1000 rpm) at 57 °C. Free cysteines were alkylated by adding iodoacetamide at a final concentration of 8.3 mM to the samples for 20 minutes in the dark. 100 µl of the samples were mixed with 50 mM Tris buffer, and then pre-digested with Trysin/Lys-C Mix (Mass spec (MS) grad, Promega). After overnight incubation at 37 °C in the dark, the samples were acidified with trifluoroacetic acid (TFA, 1% v/v). A 96 well sorbent plate (Oasis PRIME, HLB plate, Water^TM^) was equilibrated with 100% MS grade acetonitrile (ACN), and then washed with 0.1 % TFA in 99/1 H_2_O/ACN. The samples were added to each well and desalted peptides with 0.1% TFA.

### Proteomic analysis – Liquid chromatography-Tandem Mass Spectrometry (LC-MS/MS)

Peptides were reconstituted in 0.1% formic acid and applied to an Orbitrap Fusion Tribrid Mass Spectrometer with a nano-electrospray ion source as previously described. Peptides were eluted with a linear gradient of 3–8% buffer B (Acetonitrile and 0.1% Formic acid) at a flow rate of 300 nL/min over 5 minutes and then from 8–30% over a further 192 minutes. Full scans were acquired in the Orbitrap analyser using the Top Speed data dependent mode, preforming a MS scan every 3 second cycle, followed by higher energy collision-induced dissociation (HCD) MS/MS scans. MS spectra were acquired at resolution of 120,000 at 300– 1,500 m/z, RF lens 60% and an automatic gain control (AGC) ion target value of 4.0e5 for a maximum of 100 ms and an exclusion duration of 40s. MS/MS data were collected in the Ion trap using a fixed collision energy of 32% with a first mass of 110 and AGC ion target of 5.0e3 for a maximum of 100ms.

Raw data files were analysed using Peaks Studio 10.0 build 20190129. Relative protein quantification was performed using Peaks software and normalized between samples using a histone ruler ^[36]^. The mass spectrometry proteomics data have been deposited to the ProteomeXchange Consortium via the PRIDE ^[37]^.

### In vitro characterization – HBMSCs cell culture

Human bone marrow stromal cells (HBMSCs) were isolated from the human femoral bone marrow obtained from Southampton General Hospital with the approval of the local ethics committee (LREC194/99/1). The HBMSCs were isolated by repeat perfusion of the marrows and filtration through 70 µm filter before centrifugation at 1200 rpm for 5 min. The HBMSCs pellets were resuspended in minimum essential medium alpha modification (α-MEM) with supplements of 10% fetal bovine serum (FBS) and 1% penicillin/streptomycin (P/S), and culture expanded in monolayer at 37 °C and under humidified 5% CO_2_. The media was changed every 3 days.

### In vitro characterization – Cell attachment, spreading and migration

For cell attachment study, 300 µl of pre-gel solution was added to a coverslip placed in a well in 24-well culture plate. The plate was then covered with parafilm to prevent solution evaporation and incubated for 1 hour. After the incubation, HBMSCs were seeded at 1 × 10^4^ cells/well and incubated 30 minutes. They were then fixed with 2.5% glutaraldehyde in PBS. After rinsing in deionized water, the samples were dehydrated using graded ethanol (50%, 70%, 90%, 95% and 100%). The cells on the ECM hydrogels were observed using a SEM at 5 kV, and images were taken at magnifications of 55 ×, 10,000 × and 25,000 ×.

For cell migration behavior evaluation, 100 µl of pre-gel solution was dropped onto the center of a well in 12-well culture plate. The plate was then covered with parafilm to prevent the solution from evaporation and then incubated for 1 hour. While waiting for gelation, the microscope chamber was set up at 37 °C and 5% CO_2_. HBMSCs were seeded on the hydrogels in the 12-well culture plate at a density of 1× 10^4^ cells/well, and the plate was placed on the microscope stage. Time-lapse images were recorded continuously at 6 minute intervals for 24 hours.

### In vitro characterization – Cell viability and proliferation

Cell viability was investigated after 1, 3 and 7 days of culture using confocal imaging. The hydrogels were washed twice with 1× HBSS. Scaffolds were then incubated in a diluted serum-free culture media solution of Calcein AM (C3099, Invitrogen, Thermo Fisher Scientific, UK) at 37 °C and 5 % CO_2_ balanced air for 1 h, following the manufacturer’s protocol. Living cells were stained by both Calcein AM and DiD (green and red). Cells non-metabolically active (dead) were stained by DiD in red. Scaffolds were imaged using a confocal scanning microscope (Leica TCS SP5, Leica Microsystems, Wetzlar, Germany). Image analysis was carried out using Image J. Cell density was calculated normalising the number of viable cells with the volume of interest (VOI).

Cell proliferation evaluation was determined using the WST-1 colorimetric assay (Roche, Germany). ECM hydrogels were prepared in wells of 48 well plates, and HBMSCs were seeded at a density of 1× 104 cells/well and cultured for 1, 3, and 7 days. At each time point, 200 µl of WST-1 working solution was added to each well and incubated for 1 hour. The absorbance was measured at 450 nm using a microplate reader (Glomax, Promega, Southampton, UK) (n=5).

### In vitro characterization – osteogenic differentiation

#### Alkaline phosphatase (ALP) staining

Collagen and ECM gel coated surfaces (2D), and collagen and ECM hydrogels (3D) samples were prepared in 24 well plate. HBMSCs were then seeded at a density of 2 × 10^4^ cells/well with and cultured with a culture medium containing 50 mM ascorbic acid, 10 μM dexamethasone, 10 μM vitamin D3. The differentiation medium was changed every 3 days. After 7 days of cell culture, the cells were fixed in ethanol, and applying 100 μl of 4 % (v/v) Naphthol AS-MX phosphate (Sigma) and 0.0024 % fast violet B-salt (Sigma) mixed in distilled water. Cells were incubated at 37 °C for up to 60 minutes, when the reaction was stopped the images were captured and processed using a Zeiss Axiovert 200 microscope and Axiovision software V4.0. Cell Profiler software was used to calculate the relative ALP staining intensity.

DNA was quantified by the picoGreen dsDNA Quantitation Assay (Invitrogen, Paisley, UK). This assay is an ultrasensitive fluorescent nucleic acid stain of double-stranded DNA. Plates were read on an FLx cytofluor microplate reader (BioTech, Winooski, USA), and fluorescence was observed using 480 nm excitation and 520 nm emission.

Alkaline phosphatase activity was measured using a colorimetric assay (P-nitrophenol phosphate (pNPP) turnover) measuring absorbance at 410 nm on an ELx800 spectrophotometer. In brief, 10 μl of cell lysate was transferred to a 96-well clear assay plate and made up to 100 ul with 90 μl phosphatase substrate (Sigma) in 1.5 M alkaline buffer solution (Sigma). The cell lysate was incubated at 37 °C for up to 40 minutes and terminated with 100 μl of 1 M sodium hydroxide prior to reading on the spectrophotometer. ALP activity was normalized by DNA content (n=6).

#### Quantitative RT-PCR analysis

Total RNA was extracted and purified from HBMSCs that were cultured on collagen and ECM hydrogels using the Qiagen RNeasy Mini Kit. cDNA was synthesized using SuperScript VILO cDNA synthesis kit (Life Technologies, Gibco, Cambridge Bioscience). Quantitative PCR was performed using Power SYBR Green PCR Master Mix (Invitrogen Lift Technology). The reaction was made up with cDNA, SYBR Green and reverse and forward primers for the gene of interest (Table 2).

**Table 2.**
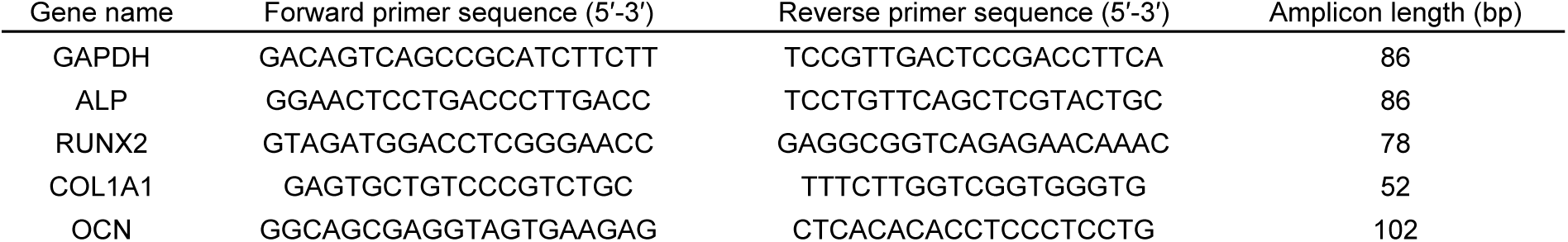
Primer sequences used for quantitative RT-PCR analysis.

All reactions were performed on the 7500 real-time PCR system (Applied Biosystems, Forster, CA) (n=4). The targeted genes expression levels were normalized to Glyceraldehyde 3-phosphate dehydrogenase (GAPDH), and were then calculated using the 2−ΔΔCt method with reference to the control group (TCP).

#### Statistical analysis

Statistical analysis was performed using GraphPad Prism 8 (GraphPad Software, La Jolla, CA) with a one-/two- ANOVA test and Tukey’s multiple comparison test. The data are presented as mean ± standard deviation (SD). Values of p < 0.05 were considered statistically significant.

## Acknowledgements

This study was supported by grants from the Biotechnology and Biological Sciences Research Council (BBSRC LO21071/ and BB/L00609X/1) and UK Regenerative Medicine Platform Hub Acellular Approaches for Therapeutic Delivery (MR/K026682/1) Acellular Hub, SMART materials 3D architecture (MR/R015651/1) and the UK Regenerative Medicine Platform (MR/L012626/1 Southampton Imaging) to RO and MRC-AMED Regenerative Medicine and Stem Cell Research Initiative (MR/ V00543X/1) to JID and YK. GC acknowledges funding from AIRC Aldi Fellowship under grant agreement No. 25412.

