## Supporting information for "Human bone tissue-derived ECM hydrogels: Controlling physicochemical, biochemical, and biological properties through processing parameters"

1. Supporting methods

***Gelation characterization – In vitro gel degradation*:** A 100 µl of pre-gel solution was dropped onto a 1.5 ml Eppendorf Protein LoBind tube and incubated for 1 hour at 37 °C incubator. Then, 1 ml of PBS was added to the tubes and incubated for 48 hours. At each time point (1, 2, 4, 8 and 24 hours), we collected 50 µl of the supernatant and directly added it into a black 96-well plate. The plate was kept at -20 °C until required. After 24 hours incubation, 400 µl of PBS containing 20 μg/ml collagenase (collagenase D, Roche) was added the tube and incubated for another 24 hours. The supernatants were collected at each time point and stored at -20 °C.

The concentration of protein in the supernatants were measured using a FluoroProfile® protein quantification kit (Sigma-Aldrich) according to the manufacture’s protocol. Briefly, 50 µl of the collected superannuates and BSA standards were added to each well of a black 96 -well plate and mixed with an equal volume of working reagent. After 30 minutes of incubation at room temperature, the fluorescent signal was measured at 520 nm (excitation) and 610 nm (emission) using a microplate reader. All the protein concentrations were normalized based on the remaining volume in the tube after collecting the supernatants at each time point.

2. Supporting figures

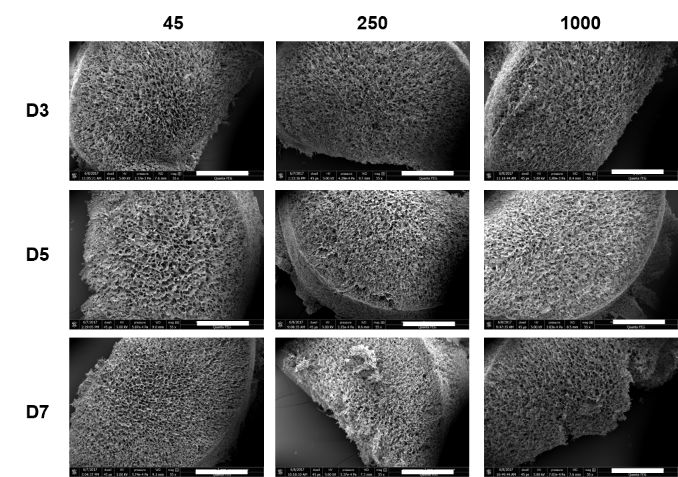

**Figure S1. Cross-section of human bone ECM hydrogels using scanning electron microscope (SEM).** All the ECM hydrogels exhibited a porous structure (scale bar = 1 mm).

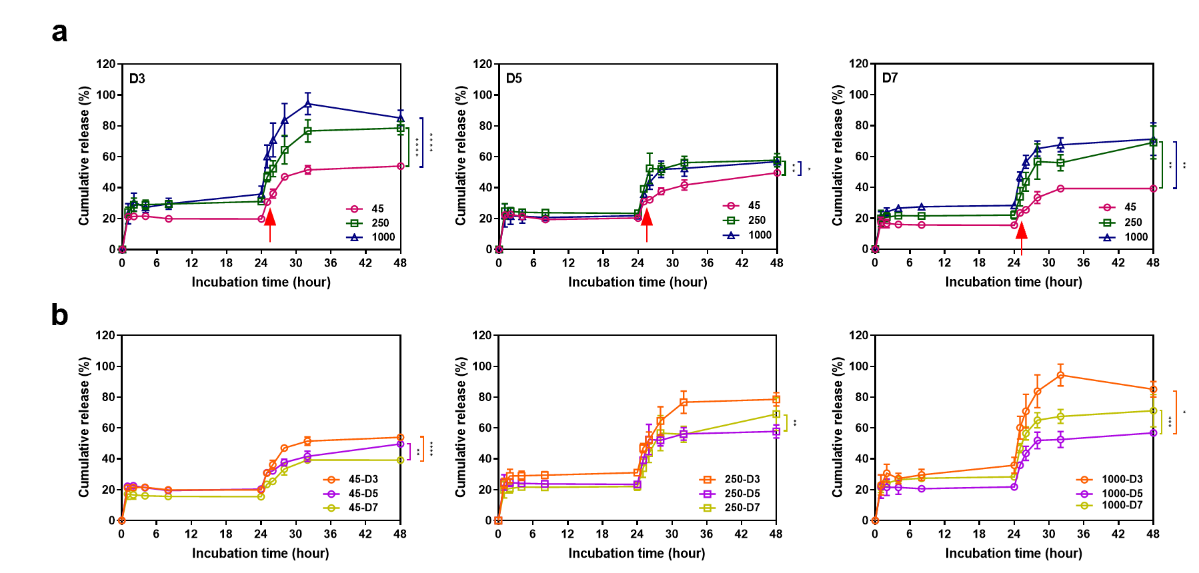

**Figure S2. Percentages of released proteins from ECM hydrogels derived from various bone powder sizes and treated for 3, 5 and 7 days.** The ECM hydrogels were incubated in PBS for 24 hours and then in collagenase PBS for another 24 hours (indicated with arrows), followed by measuring protein concentrations. The bone powder size and digestion time significantly affected the degradation of ECM hydrogels. Statistical analysis was performed using one-way ANOVA followed by Tukey’s multiple comparisons test (*p<0.05 **p<0.005 ***p<0.0005 ****p<0.0001). Data represent mean ± SD, N = 4.

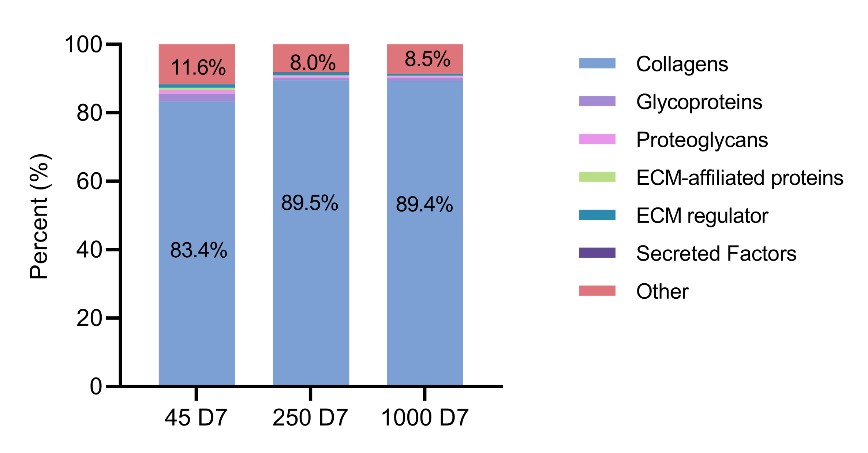

**Figure S3. Comparison of the proportion of subcategorized proteins in human bone ECM digests derived from various bone powder sizes and treated for 7 days.** The ECM hydrogels derived from 45 – 250 µm size of bone powders and treated in pepsin solution for 7 days induced a variety of ECM proteins.
